# Relationship between shear wave velocity and muscle activation is inconsistent across different muscle types

**DOI:** 10.1101/2021.04.07.438761

**Authors:** Michel Bernabei, Daniel Ludvig, Thomas G. Sandercock, Eric J. Perreault, Sabrina S. M. Lee

## Abstract

There is an increasing use of shear wave ultrasound elastography to quantify mechanical properties of muscles under various conditions such as changes muscle length and levels of activation in healthy and pathological muscle. However, little is known about the variability in shear wave velocity among muscles as most studies investigate one specific muscle. The purpose of this study was to determine if the relationship between SWV and muscle activation is consistent across muscles with different architectures: biceps brachii, tibialis anterior, and medial gastrocnemius, All measures were made at matching levels of activation and approximately at the optimal length for each muscle to control for length-dependent changes in the relationship between activation and force or stiffness. We also conducted a control experiment to determine how the passive force within a muscle alters the relationship between muscle activation and shear wave velocity. The relationship between shear wave velocity-squared and activation above 10% MVC differed across muscles, with biceps brachii and medial gastrocnemius showing a lower slope than tibialis anterior. Shear wave velocity-squared also differed between muscles at the shortest length (p<0.001) and the increase in shear wave velocity-squared with muscle lengthening also differed between muscle types (p = 0.005) Muscle-specific differences could not be explained by the architectural features such as pennation angle, fiber length, and physiological cross-sectional area. Our results demonstrate that there is not a unique relationship between muscle activation and shear wave velocity, highlighting the importance of understanding the many factors contributing to shear wave propagation in muscle before elastography can be used to make quantitative comparisons across muscle types.

## INTRODUCTION

The potential for evaluating changes in muscle properties associated with various pathologies has driven much ultrasound development. B-mode ultrasound has proven to be useful for assessing a variety of architectural parameters such as fascicle lengths and pennation angles in healthy individuals and in those with impairments including from cerebral palsy or stroke. Ultrasound shear wave elastography has emerged as a potential tool for estimating the mechanical properties muscle in healthy (Chernak et al., 2013; Maïsetti et al., 2012; Nordez and Hug, 2010) and diseased states (Abel et al., 2003; Lacourpaille et al., 2014; Lee et al., 2015), and is often used under the assumption that shear wave velocity (SWV) varies in proportion to the stiffness of a muscle (Alfuraih et al., 2019; Du et al., 2016; Eby et al., 2015). However, is remains unknown if the relationship between SWV and stiffness is consistent across muscles.

Many studies have demonstrated that SWV changes under conditions that also change muscle stiffness. It varies almost linearly with muscle activation (Chernak et al., 2013; Lapole et al., 2015; Nordez and Hug, 2010; Yoshitake et al., 2013) and with passive stretching (Koo et al., 2014; Le Sant et al., 2017). Several pathological conditions known to increase muscle stiffness also exhibit increased SWV, including stroke (Le Sant et al., 2019; Lee et al., 2015) and cerebral palsy (Kwon, 2012; Lee et al., 2016). While these studies have demonstrated that SWV can reliably detect chances in muscle stiffness, it has not been established if there is a unique relationship between these variables, as would be needed to quantify muscle stiffness across different muscles or individuals with different physiological states.

It has long been known that muscle stiffness increases with muscle stress. This increase is consistent across muscles with very different architectures provided that fascicle lengths are considered (Cui et al., 2008). These results arise from an activation-dependent change in the Young’s modulus of muscle, presumably resulting from an activation-dependent change in the number of active cross-bridges. If SWV can be used to provide a consistent, quantitative measure of muscle stiffness, it too should scale with muscle activation in manner that is consistent across muscles with different architectures.

The purpose of this study was to determine if the relationship between SWV and muscle activation is consistent across muscles with different architectures. We made measurements in three muscles that covered a range of pennation angles, fascicle lengths and physiological cross-sectional areas. We hypothesized that the dependency of SWV on activation would be consistent across muscles. All measures were made approximately at the optimal length for each muscle to control for length-dependent changes in the relationship between activation and force or stiffness. We also conducted a control experiment to determine how the passive force within a muscle alters the relationship between muscle activation and SWV. Together, our results characterize the variability that can be expected when using ultrasound shear wave elastography to estimate the stiffness of individual muscles.

## METHODS

We designed our experiments to determine if there is a consistent relationship between muscle activation and SWV in muscles with highly differing architectures. Two experiments were conducted. The primary experiment assessed the SWV-activation relationship near the optimal length for each muscle. This was done to control for length-dependent changes in the activation force, and therefore stiffness, relationship for each muscle. The second experiment was conducted at the shortest length that could be obtained for each muscle. This was to mitigate the effects of passive tension on SWV (Koo et al., 2013; Le Sant et al., 2017; Martin et al., 2018) which would likely be different between muscles when studied only at their optimal length. Only passive measurements were made at the shortest length.

### Subjects

Twelve healthy young adults took part in this study (7 females and 5 males, mean (SD) age: 29 (5) years; height: 1.72 (0.11) m; body mass: 73.7 (14.2) kg). Subjects had no history of major orthopedic diagnosis, musculoskeletal trauma, or persistent joint pain at the targeted body locations. All experimental procedures were approved by Northwestern University’s Institutional Review Board (#0020-33-96), in compliance with the Helsinki Declaration. All subjects received and signed informed consent prior to collecting data.

### Setup

The primary goals of this study were to assess the relationship between SWV and activation, as well the relationship between SWV and muscle length under passive conditions, across muscles with different architecture. We chose three muscles: i) the long head of the biceps brachii (BB), ii) the medial gastrocnemius (MG), and iii) the tibialis anterior (TA) because of their remarkably different architectural features (Table 1). Participants sat upright on a Biodex (Biodex Medical Systems Inc., Shirley, USA) chair with the hip at 90 degrees flexion and knee at the most extended position possible (0-10 degrees flexion). Their dominant forearm or lower leg was secured in a neutral position with a fiberglass cast. The cast was then secured to a metal plate attached to the shaft of the Biodex motor and aligned to its rotational axis (Fig. 1A). The Biodex motor was set to position control, maintaining the relevant joint at the desired angle. Joint torques were measured with a six degrees-of-freedom load cell (JR3 Inc., Woodland, USA), attached to the Biodex motor shaft. Surface EMG (Bagnoli EMG system, Delsys Inc., Boston, MA; 20-450Hz bandpass) was recorded using bipolar electrodes (#272, Noraxon, USA) placed on the targeted muscles as well as synergistic and antagonistic muscles (triceps brachii, MG, and soleus) to account for effects of co-activation. EMG, load cell and ultrasound trigger signals were sampled (2 kHz) and digitized with a 16-bit data acquisition board (USB-6343, National Instruments, Austin, US).

**Table 1.**
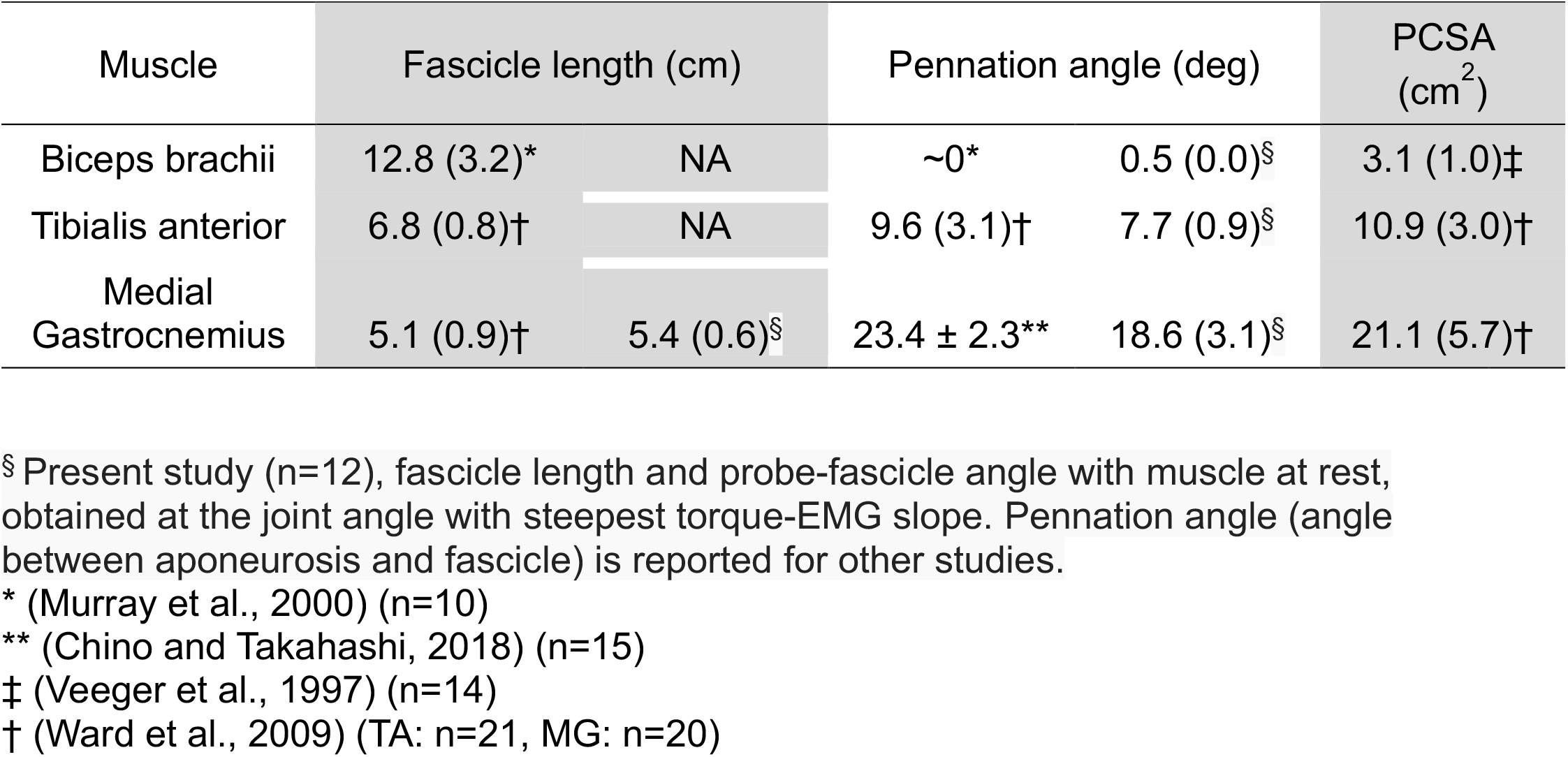
Architectural characteristics of target muscles | mean (SD)

| Muscle | Fascicle length (cm) |  | Pennation angle (deg) |  | PCSA (cm<sup>2</sup>) |
| --- | --- | --- | --- | --- | --- |
| Biceps brachii | 12.8 (3.2)* | NA | ~0* | 0.5 (0.0)§ | 3.1 (1.0)‡ |
| Tibialis anterior | 6.8 (0.8)† | NA | 9.6 (3.1)† | 7.7 (0.9)§ | 10.9 (3.0)† |
| Medial Gastrocnemius | 5.1 (0.9)† | 5.4 (0.6)§ | 23.4 ± 2.3** | 18.6 (3.1)§ | 21.1 (5.7)† |

§ Present study (n=12), fascicle length and probe-fascicle angle with muscle at rest, obtained at the joint angle with steepest torque-EMG slope. Pennation angle (angle between aponeurosis and fascicle) is reported for other studies.
\* (Murray et al., 2000) (n=10)
\*\* (Chino and Takahashi, 2018) (n=15)
‡ (Veeger et al., 1997) (n=14)
† (Ward et al., 2009) (TA: n=21, MG: n=20)

**Figure 1.**
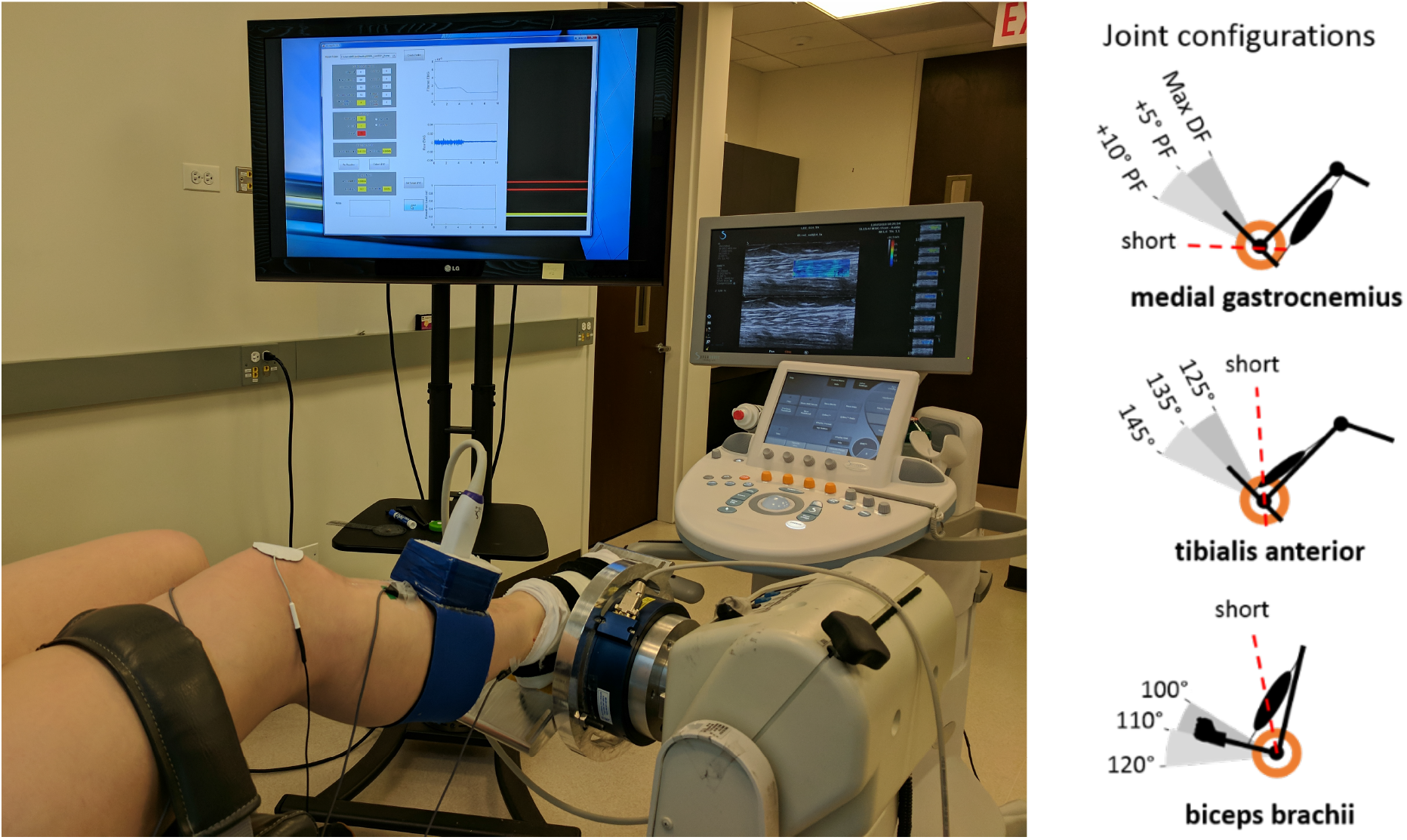
Experimental setup for measuring shear wave velocity (SWV), torque and activation of the tibialis anterior. The subject was secured to a Biodex chair by a strap on the thigh. The foot was cast and secured to a rotating foot plate by Velcro straps and connected to the load cell. Bipolar surface electrodes were placed on the target muscle (tibialis anterior) and its main antagonists (medial and later gastrocnemii). The ultrasound transducer was positioned on the mid-belly of the muscle. B) Different joint angle configurations were imposed for each of the three muscles, based on their specific physiological range of motion. An additional ‘short’ joint configuration corresponded to the smallest joint angle achievable in experimental conditions; this condition minimizes the effect of passive tension on shear wave measures. BB: biceps brachii; TB: triceps brachii; DF: dorsiflexion; PF: plantarflexion.

### Ultrasound measurements

The ultrasound transducer was secured on the muscle of interest with a neoprene custom-molded holder and positioned at the mid-belly region of BB and at the proximal region of MG and TA muscle bellies to avoid internal tendons within each muscle. The imaging plane was aligned with the fascicle plane as viewed from the B-mode image. The region for elastography was a 25×10 mm rectangle between the superficial and deep aponeuroses (Fig. 2A). For each isometric contraction a B-mode ultrasound image and a SW propagation video were simultaneously captured using a research grade ultrasonography system (Aixplorer SuperSonic Imagine, Aix-en-Provence, France) with a linear transducer array (4-15 MHz, SuperLinear 15-4, Vermon, France). The parameters of the system were set to i) mode: MSK – foot and ankle ii) opt: std iii) persist: no, to maximize the sampling rate and avoid bias due to locked proprietary data post-processing. The software on our system provides SW movie data sampled at 8 kHz in sequences of 42 frames for each acoustic radiation pulse, recorded simultaneously with a single B-mode ultrasound image.

**Figure 2.**
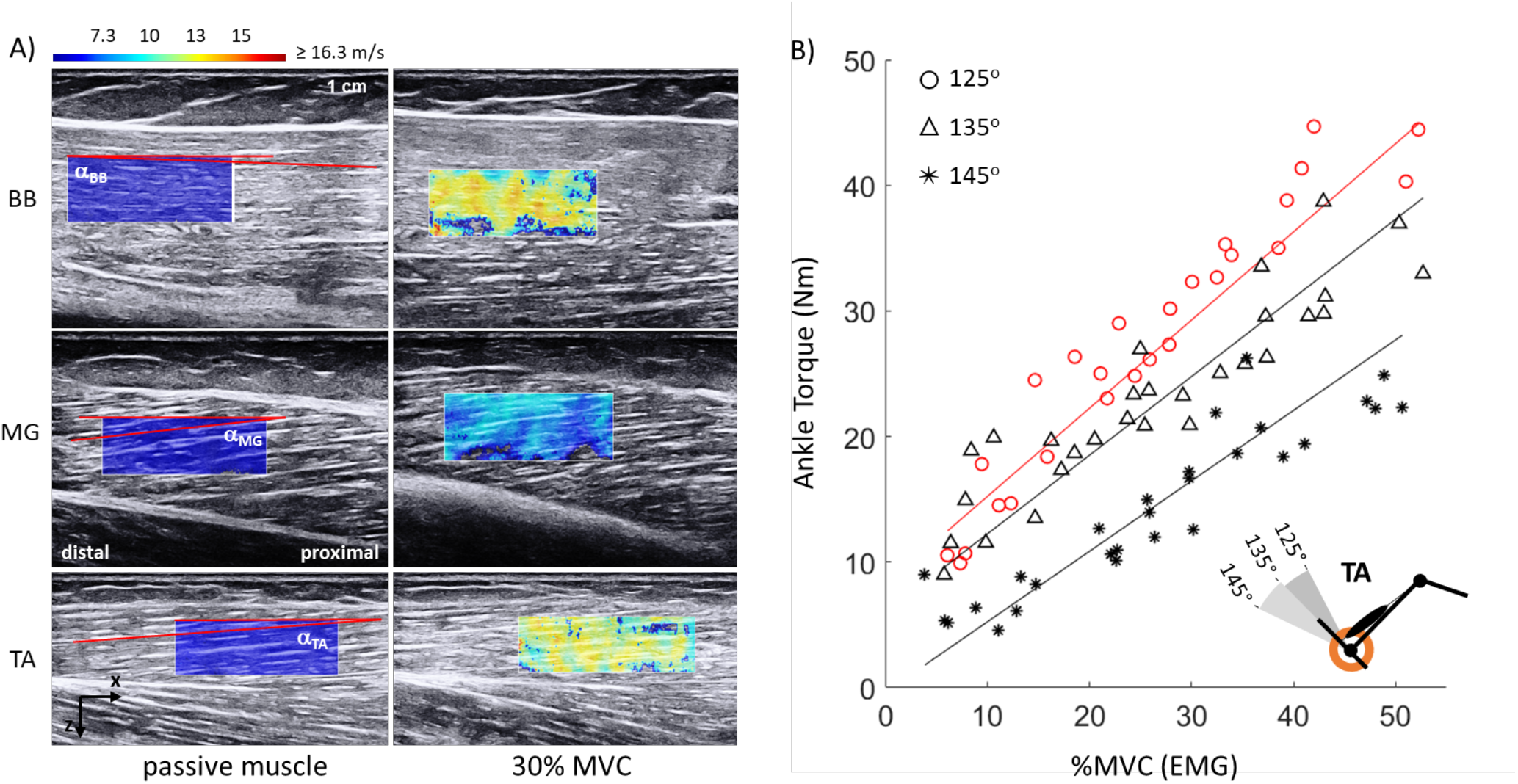
Ultrasound and torque data with different joint angle configurations and muscle activations. A) Heatmaps representing local shear wave velocity estimates on biceps brachii (BB), medial gastrocnemius (MG) and tibialis anterior (TA) for passive (left panels) and active muscle state (30% MVC, right panels). Red lines show an example of probe-fascicle angle measures (α) extracted for each muscle and condition. Note that MG heatmap shows qualitatively lower shear wave velocity, as calculated by the AIXplorer, compared to BB and TA at matching level of activation (30% MVC). B) Exemplar data of ankle torque measured in TA with increasing levels of activation and at three different joint angle configurations. Note that ankle torque covaried with muscle activation and its rate of change was different across ankle angles. The angle at which the highest slope occurred was used to estimate optimal length (red).

### Protocol

In the primary experiment, we assessed the relationship between SWV and muscle activation at three different lengths for each muscle by changing the joint angle. The three angles were selected to position the muscle near its optimal length, based on values reported in the literature. The selected angles were 100-110-120° for BB (Leedham and Dowling, 1995), 35-45-55° for TA (Maganaris, 2001); maximum dorsiflexion to +5° and +10° plantarflexion for MG (Fig. 1B) (Maganaris, 2003). Each muscle was tested in a separate session.

For each muscle at each joint angle, passive and maximum voluntary contraction (MVC) trials were performed at the beginning of each session to normalize EMG data. Trials were then conducted at increasing levels of muscle activation from 0 to 30% MVC in 5% MVC steps, and at 40, 50, and 75% MVC. EMG feedback was provided to the subject to assist with task completion. The EMG feedback signal was digitally rectified and low-pass filtered (1Hz low-pass, II-order Bessel) before being displayed to the subject. At each imposed joint angle, three trials (five seconds each) at each level of activation were conducted. Trials were randomized across levels of muscle activation and imposed joint angles. A total of 90 trials were conducted for each muscle, excluding baseline and MVC trials.

The second experiment involved making measurements at the shortest possible muscle length and was conducted on a separate day (all muscles) than the first experiment. A subset of the subjects (n = 9) from the first experiment participated in this second experiment. Subjects were seated as described above for the first experiment. For each muscle, the corresponding joint was moved to a position where the muscle would be at its shortest length possible while still being able to position the ultrasound transducer to capture images (SWV elastography and Bmode). Three trials were conducted for each muscle with the order of muscle randomized. Measurements were only made under passive conditions.

### Data analysis

All EMG data were detrended and rectified before further processing. The mean EMGs recorded during the MVCs for each experimental condition were used to normalize the EMGs recorded during all submaximal contractions. The muscle activations and joint torques corresponding to each ultrasound measurement were estimated by averaging the EMG and torque recordings over a period of 450 ms immediately prior to the start of ultrasound imaging.

To create an equitable comparison of the SWV-EMG relationships across all muscles, we attempted to place each muscle as close as possible to its optimal length. For each of the three joint angles tested, we found the slope of the EMG-torque relationship (Fig. 2B). We chose the angle with the highest slope as the angle where the muscle was closest to its optimal length. Only data from this joint angle was used for further analysis.

Shear wave ultrasound data were processed using a custom-written MATLAB program where SWV was estimated using a time-to-peak method (Bernabei et al., 2020). Most of our results are reported in terms of SWV^2^ since changes in muscle stiffness are expected to vary in proportion to the square of SWV (Bercoff et al., 2004; Royer et al., 2011).

In each position, probe-fascicle angle and fascicle length were extracted from ultrasonic images of the targeted muscles. Typically, pennation angle, the angle between the fascicle and aponeurosis, is measured as an architectural parameter. In this study, we quantify probe-fascicle angle instead, since this is most relevant to our method for inducing shear waves in muscle and measuring their propagation. The probe-fascicle angle (α) was defined as the angle between a fascicle and the horizontal line of the region of interest for elastography provided by the Aixplorer in the DICOM image (Fig. 2A, color map rectangle). This angle is useful to assess if fascicle orientation affects shear wave propagation as the shear waves travel parallel to the probe direction; thus, the horizontal line of the region interest is used for reference. Fascicle length was estimated by tracing the same fascicle from the superficial aponeurosis to the deep one. Images were pre-processed to enhance contrast of the features of interest. All architectural measurements were completed by a single operator; the known reproducibility of this measure (Kawakami et al., 1998) was confirmed here by a correlation coefficient of 0.997 across three repeated measurement sessions over subsequent days, on all probe-fascicle angle data from the same muscle (30 samples x 3 days).

### Statistics

Linear mixed-effects models were used for all hypothesis tests. Our primary hypothesis was that the SWV-muscle activation relationship is consistent across muscle types. We tested this with the squared SWV as the dependent factor; EMG amplitude as a continuous factor, the three muscles as fixed factors, and each subject as a random factor. In addition to our primary hypothesis, we tested two additional hypotheses. The first was that SWV would be consistent when the muscle is at a very short length. We tested this with the squared SWV as dependent factor, muscle as a fixed factor and subject as a random factor. Post-hoc ANOVAs and Wald t-statistics with Satterthwaite corrections were used to test model significance and the overall significance of each modeled predictor. Significance was assessed at the p < 0.05 level and fixed effects estimates reported as average values with 95% confidence interval. All statistical tests, as well as all signal and image processing, were performed in MATLAB (Mathworks, Natick, USA).

## RESULTS

### Effects of activation on shear velocity across muscles

The relationship between muscle activation and shear-wave velocity differed drastically between muscle types. Shear wave velocity-squared increased with muscle activation as can be seen in the representative subject shown in Fig. 3A. This increase in SWV^2^ with muscle activation happened in two stages: a passive value and initial linear increase levels to ∼10% MVC; and a second linear increase with a different slope from 10% MVC to 50% MVC. Here we focus on the latter phase covering the broadest range of activation levels; the passive measurements are considered in the following section. The relationship between SWV^2^ and activation above 10% MVC differed across muscles, with BB and MG showing a lower slope than TA. For all subjects, the linear mixed-effects model (Fig. 3B) accurately modeled the data (R^2^ = 0.75). Contrary to our null hypothesis, there was a significant difference between the SWV^2^ to EMG relationships measured across muscles (F_2,36_ = 12.4, p<0.001). The slope of this relationship was significantly greater in TA (5.0 ± 0.3 (m^2^/s^2^)*(%MVC)^-1^) than in the MG (2.9 ± 0.3 (m^2^/s^2^)*(%MVC)-^1^, t_35_ = 4.2, p<0.001) or the BB (2.9 ± 0.3, (m^2^/s^2^)*(%MVC)^-1^, t_36_ = 4.5, p<0.001). There was little difference in the slope between MG and BB (t_36_ = 0.1, p = 0.90).

**Figure 3.**
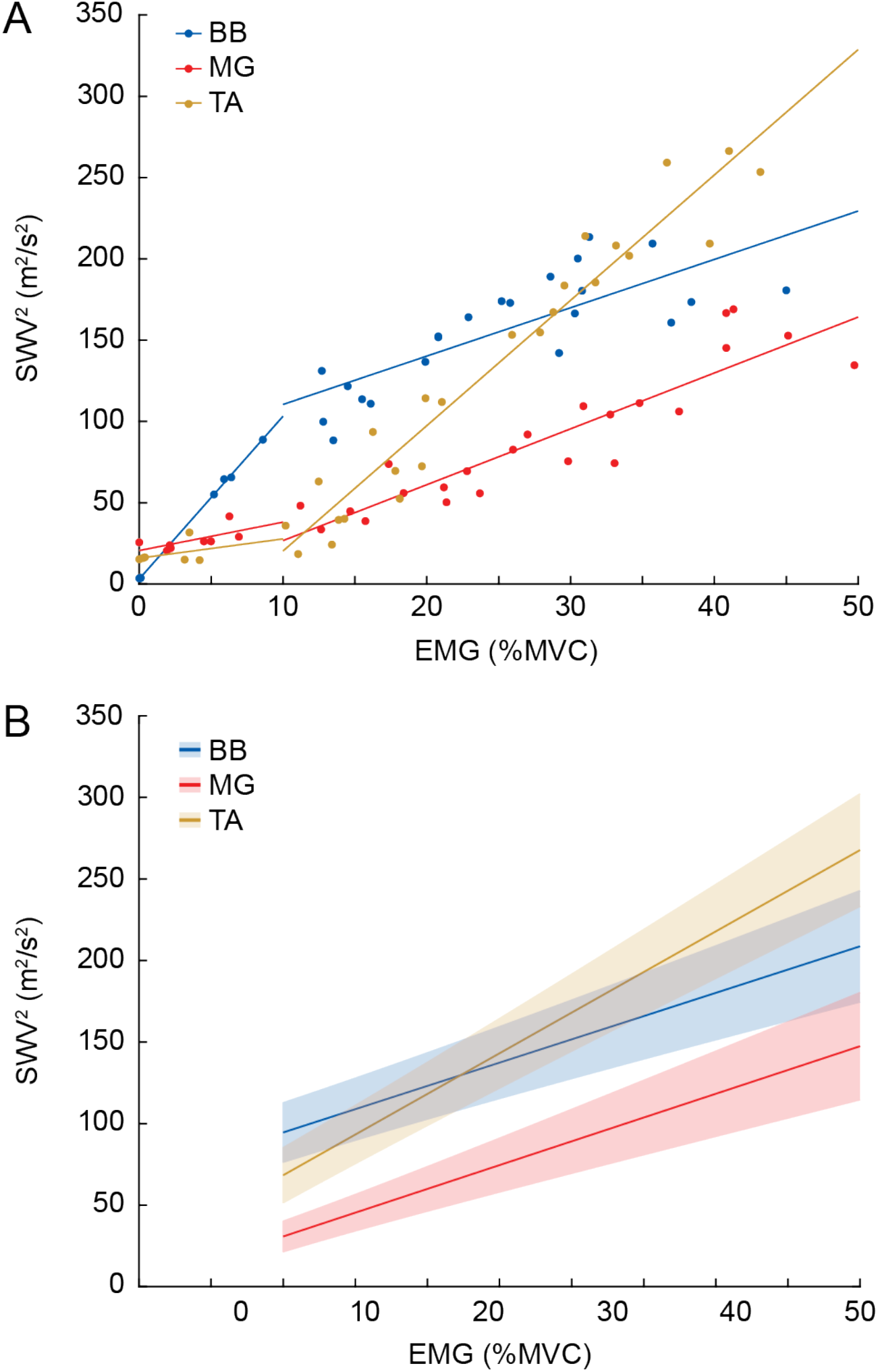
The relationship between shear wave velocity and isometric contraction level differed between muscles. A) Data for an example subject showing how shear wave velocity squared (SWV^2^) increased linearly with increasing muscle contraction as measured by EMG. B) Group results showing how the slope of the shear wave velocity squared-EMG slope was greatest in the tibialis anterior (yellow) compared to the biceps brachii (blue) and medial gastrocnemius (red). The solid lines indicate the estimated shear wave velocity-squared from the linear mixed-effects model, with the shaded region representing the 95% confidence intervals.

### Effects of muscle length on shear wave velocity during passive conditions

One potential explanation for the activation-dependent differences between muscles shown in Fig. 3 is that muscles had significantly different passive stresses at optimal length, where the primary experiment was conducted. For this reason, we also assessed the length-dependent change in passive SWV at the shortest attainable lengths for each muscle, and at the three lengths used in the primary experiment.

The relationship between SWV^2^ and muscle length during passive lengthening differed between muscle types. The values of SWV^2^ were substantially different across muscles in resting conditions even at their shortest lengths (Fig. 4A, B, and C; ‘short’, F_1,16_ = 18.5, p < 0.001), where we expected differences due to passive muscle tension to be minimal. The BB had the lowest SWV^2^ (3.1 ± 1.7 m^2^/s^2^), significantly lower than MG (5.5 ± 1.7 m^2^/s^2^, t_16_=2.1, p=.05) and the TA (9.8 ± 1.7 m^2^/s^2^, t_16_=6.0, p<0.001). Shear wave velocity-squared in the MG was also significantly lower than the SWV^2^ in the TA (t_16_=4.0, p=.0012). When changing joint angle to passively stretch each target muscle to regions reported to be near optimal length (see Methods), measures of SWV^2^ increased from the shorter to longer muscle lengths (average increase 0.76 ± 0.15 m^2^/(s^2^*deg). This increase in SWV^2^ with muscle lengthening did differ between muscle types (Figure 4D; F_2,79_=58, p<0.001). The increase was significantly smaller in the BB (0.13 ± 0.21 m^2^/(s^2^*deg)), compared to the MG (1.59 ± 0.34 m^2^/(s^2^*deg), t_87_=7.3, p<0.001) or TA (0.55 ± 0.21 m^2^/(s^2^*deg), t_66_=2.9, p = 0.005). This increase was also significantly larger in the MG than the TA (t_87_=5.2, p<0.001). This demonstrates that even under passive condition SWV^2^ is not consistent across muscle types.

**Figure 4.**
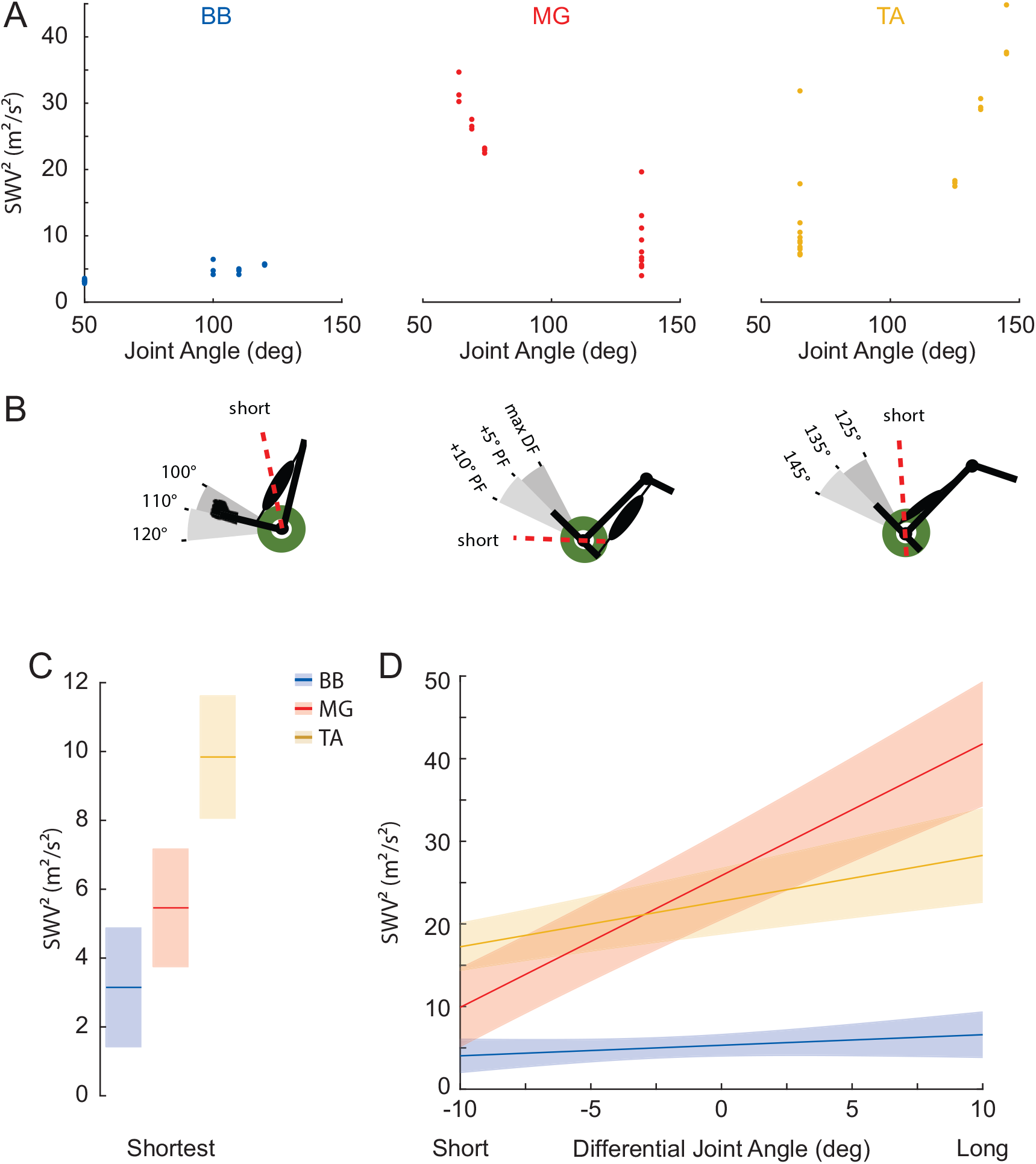
Dependency of shear wave velocity on passive muscle length across muscle types. A) Shear wave velocity squared (SWV^2^) increased with lengthening the target muscles as shown for a representative subject. B) Joint configurations for the three muscles tested. We measured SWV^2^ at a joint angle that resulted in the shortest possible muscle length (BB: ∼40°; MG: max PF; TA: max DF), as well as 3 angles, 10 degrees apart, centered near each muscle’s optimal length. C) In the shortest muscle condition, the SWV^2^ was greatest in the TA and smallest in the BB. D) The MG showed the greatest increase with passive lengthening, followed by the TA; there was only a slight increase in SWV^2^ when lengthening in the BB.

## DISCUSSION

This study examined if SWV was different across three muscles with different architectural features, including pennation angle, fascicle length, and PCSA. We found that the relationship between SWV and activation was different across the BB, TA, and MG. Even at very short lengths under passive conditions where muscle force is expected to be at a minimum, SWV was different across the three muscles. None of the architectural features we considered could explain the observed findings since the TA, which had architectural features between the other two muscles (Table 1), consistently had the steepest change in SWV with activation (Fig. 3). These results emphasize the need to have a more complete understanding of the factors contributing to how shear waves propagate in muscle before the promise of elastography can be used to make intermuscular comparisons for muscles with highly divergent architectures.

Our primary finding was that the relationship between muscle activation and SWV differed across muscle types. While there have been many studies on how SWV changes with muscle activation (Chernak et al., 2013; Lapole et al., 2015; Nordez and Hug, 2010; Yoshitake et al., 2013), our work is the first to determine the consistency of this relationship across muscles with different architectures. The range of SWV values collected from 5 to 50% MVC, were much lower in MG compared to BB and TA (Fig. 3) and the rate of increase varied drastically between BB, MG, and TA which had the greatest slope. One of the main challenges of comparing SWV across muscles was to exclude the possibility that the variability is a consequence of differences in the EMG-torque relationship specific to each muscle. Thus, we compared SWV across muscle lengths near the optimal range that corresponded to the maximum rate of change for joint torque. Without knowing the optimal length of the muscles, we chose the length, modified by joint angle, at which the slope of the SWV^2^ – activation was the greatest. An approach to determine SWV over the full range of muscle length under both passive and active conditions across muscles would allow a more comprehensive assessment of how SWV varies across muscles given that SWV is sensitive to both activation and muscle length.

SWV is sensitive to the passive stress in muscle (Bernabei, 2019; Le Sant et al., 2017). We therefore explored how passive lengthening changed SWV in each of our studied muscles. The relationship between SWV and passive lengthening is well documented for individual muscles (Le Sant et al., 2017; Maïsetti et al., 2012), but comparison across muscles is limited (Le Sant et al., 2017). Shear wave velocity was substantially different across muscles (Fig. 4) even at the shortest length possible (for which we could place the ultrasound probe) and where there was minimal joint torque. When passively stretched, the increase in SWV also varied across the muscles, demonstrating that even under passive conditions, SWV is not consistent across muscle types. This is similar to observations of differences in SWV for several lower extremity muscles of the ankle during passive lengthening by Le Sant el al. (2017). They attributed these differences in SWV to variations in muscle properties which influence passive stress such as passive force-length relationship, physiological cross-sectional area, and moment arms.

Our results could not be explained by the architectural differences we considered including pennation angle, fascicle length, and PCSA; the relationship between muscle activation and SWV was not affected by variations in muscle architecture (Table 1). We also considered the probe-fascicle angle which slightly differs from the pennation angle; instead of measuring the fascicle orientation relative to the aponeurois as commonly done in the literature, the angle between fascicles and the horizontal direction of the probe which is parallel to the shear wave propagation was considered; however, no effect was found. This is supported by observations of only small effects (<1.3 % for an angle of max 20°; (Miyamoto et al., 2015)) of tilting the probe within the plane of the fascicles when imaging the parallel-fiber BB. It is possible that variations in three-dimensional fascicle orientation such as out-of-plane fascicle curvature across muscles may affect SWV measures (Bouchet et al., 2020; Eby et al., 2013). These results suggests that other muscle properties not explored in this study may influence SWV.

Factors we were unable to measure may also have contributed to our results. In particular, composition may influence shear wave propagation. There are differences in the collagen content of the BB, TA, and MG (10.0 mg/mg, 10.7 mg/mg, and 13.4 mg/mg; (Lin, 2011) though the effect on SWV requires further investigation. We have previously demonstrated that SWV is sensitive to passive stress in muscle in an animal model (Bernabei, 2019). As direct measurements of stress are not possible within muscle, local variations in strain may be the most informative about stress. Local variations in strain (Blemker et al., 2005; Jensen et al., 2016) occur in muscles with complex internal architectures such as internal tendons of the TA and BB, respectively, under passive and active conditions. These non-uniform strains may lead to non-uniform stresses within the muscle such that depending on where SWV is measured, SWV values will vary.

## CONCLUSIONS

Our results demonstrate how SWV sensitivity to both changes in muscle activation and passive lengthening differ across muscle types. These results could not be explained by the architectural features of muscle that we measured, but could result from unmeasured differences in material composition or localized differences in the internal stress of each muscle. Exploring cause of our observed differences will be helpful for extending the use of muscle shear wave elastography into a quantitative tool that can be used to assess muscle properties across a range of muscle types and physiological conditions.

## ACKNOWLEDGMENTS

The authors thank Allison Bingqing Wang and Kristen Jakubowski for the preliminary analysis of ultrasound elastography images.

## AUTHOR CONTRIBUTIONS

S.S.M.L., E.J.P., and T.G.S. conceived and designed research; M.B. and S.S.M.L. performed experiments; M.B. and D.L. analyzed data; E.J.P., M.B., S.S.M.L., D.L., and T.G.S. interpreted results of experiments; M.B. and D.L. prepared figures; M.B. drafted manuscript; E.J.P., M.B., S.S.M.L., D.L., and T.G.S. edited and revised manuscript; M.B., S.S.M.L., D.L., E.J.P., and T.G.S. approved final version of manuscript.

